# Initiation codon context governs translation-coupled mRNA decay and coordinated expression in the human parasite *Leishmania*

**DOI:** 10.64898/2026.07.01.735800

**Authors:** Ana Maria M. Santi, Thomas Cokelaer, Juliana Pipoli da Fonseca, Paul Jenkins, Elena B. Tikhonova, Cristian C. Rodriguez-Almonacid, Andrey L. Karamyshev, Zemfira N. Karamysheva, Gerald F. Späth

## Abstract

In the absence of canonical transcriptional regulation, *Leishmania* expression changes largely rely on post-transcriptional control via differential mRNA turnover, yet how this process is linked to translation-dependent decay remains poorly understood. Here, by using the initiation codon sequence context as a molecular entry point, we uncover that Kozak-mediated changes in translational efficiency affect mRNA stability. Genome-wide mapping of Kozak sequences across pathogenic trypanosomatids revealed evolutionary adaptation of this *cis*-regulatory element at the genus and sub-genus levels, as judged by the selective enrichment of only a small subset of possible Kozak motifs that clustered in a species-specific fashion. Transgenic expression of EGFP under the control of distinct Kozak variants correlated EGFP mRNA recruitment to monosomes and heavy polysomes with respectively decreased and increased protein abundance. Surprisingly, this correlation also extended to EGFP transcript abundance, suggesting a role of the Kozak motif in translation-mediated mRNA turnover. Transcript profiling of parasites arrested for either transcription or translation elongation revealed a subset of unstable mRNAs that were rescued from decay in a translation-dependent fashion. Significantly, these transcripts contained a characteristic Kozak motif further supporting a possible role of the initiation codon sequence context in co-regulation of translation-mediated mRNA decay. Our work uncovers a key role for translation initiation-coupled mRNA decay in *Leishmania* gene expression regulation, adding a previously under-appreciated layer of post-transcriptional regulation in parasite adaptation.

## INTRODUCTION

The leishmaniases are a complex of parasitic diseases caused by various species of the protozoan genus *Leishmania*, which are transmitted to mammalian hosts by female phlebotomine sand flies ^1^. With an endemic footprint spanning 90 countries and territories, leishmaniasis represents a substantial global health challenge. It ranks among the most lethal neglected tropical diseases and primarily afflicts impoverished populations, highlighting its role as both a public health and socioeconomic threat ^1^.

Despite the absence of promoter-driven gene regulation*, Leishmania* parasites exhibit remarkable phenotypic plasticity and adaptability ^2^. This is linked to their atypical genome architecture, in which nearly all protein-coding genes lack introns and are arranged into long, unidirectional polycistronic clusters without overt functional linkage ^3–7^. These polycistronic pre-mRNAs are processed into individual mature transcripts ^7,8^. Although transcription is broadly constitutive and mediated by RNA polymerase II, the mechanisms of transcription initiation remain poorly defined due to the lack of identifiable canonical promoters ^4,9^. Emerging evidence suggests that epigenetic factors modulate DNA accessibility to influence transcription start sites ^10–15^. Termination, in contrast, is directed by the modified nucleotide base J (β-D-glucosyl-hydroxymethyluracil), a thymine modification found at the end of each polycistronic unit ^16–19^.

In compensation for the lack of transcriptional regulation, gene expression in *Leishmania* is predominantly controlled at the epigenomic and post-transcriptional level. While chromosomal copy number variation can influence transcript abundance ^20–23^, transcript and protein levels are frequently uncorrelated, underscoring the importance of downstream regulation ^24–26^. Post-transcriptional regulation in *Leishmania* operates through mechanisms that govern stability, processing, transport, degradation, and translational efficiency of mRNA, or protein turnover ^20–29^. At the level of translation, mRNAs differentially compete for ribosomal subunits based on their structural elements, codon bias, modifications, and presence of RNA-binding proteins (RBPs) or regulatory ncRNAs, thereby selectively influencing protein production rates ^28–34^. Ribosome heterogeneity itself, arising from variations in rRNA modification, ribosomal protein composition and post-translational modifications, can be dynamically modulated by developmental or environmental signals, adding an additional layer of regulatory complexity ^35–38^. Indeed, we have previously demonstrated that *L. donovani* achieves fitness gain during culture adaptation through frequent changes in chromosome and gene copy numbers, with deleterious gene dosage effects being buffered at post-transcriptional levels, thus establishing stable fitness phenotypes ^38^. The same processes regulating stage-specific gene expression levels were also shown by us and others in recent multi-layer systems analyses ^37,39^.

An important step of translational control occurs at initiation, when mRNAs are recruited to ribosomes ^40^. The 5’ untranslated region (5’UTR) contains multiple regulatory elements that influence translation efficiency in eukaryotes, including its length ^41^, RNA secondary structures, AU-rich motifs, upstream open reading frames (uORFs), GC content, polypurine tracts, internal ribosome entry sites (IRES), 5’ cap structure, and translation enhancers ^42,43^. Most importantly, the nucleotide context surrounding the start codon (AUG), known as the Kozak sequence, influences the probability of ribosome recognition during the scanning process at the initiation of translation ^43–45^. Despite the documented role of Kozak sequences in regulating translation across diverse eukaryotes, their functional impact in trypanosomatids remains largely unexplored. While early studies in *Leishmania* demonstrated that these sequences can modulate translation of reporter genes in *L. tropica* and *L. tarentolae* ^46,47^, to our knowledge, no further investigation has been conducted in the two decades since.

Here, by combining computational, genetic and pharmacological approaches with transcript profiling, our study reveals an important role of the *Leishmania* Kozak sequence (defined here as positions -5 to -1 relative to the start codon) in gene expression by coordinating translation-mediated mRNA turnover. This process may play an important role in *Leishmania* differential gene expression during parasite life cycle progression, and in harnessing genome instability for fitness gain through the compensation of toxic gene dosage effects.

## MATERIALS AND METHODS

### Kozak Sequence Extraction and analysis

Kozak sequences were extracted from different genomes using the open-source KozakExplorer pipeline ^48^ (https://github.com/sequana/webapp_kozak), which is part of the Sequana project ^49^.

For t-SNE analysis, we used Kozak Explorer for a broad comparative analysis across the tree of life. Seventy-two euglenozoan genomes spanning multiple kinetoplastid genera were retrieved from NCBI GenBank v1.0.0 (Supplementary Table 1). For each genome, Kozak sequence profiles were extracted from the analysis window (−20 to +6 positions relative to the AUG start codon, AUG excluded) using the sequana.kozak module. Feature vectors for dimensionality reduction were constructed by concatenating per-position Kullback-Leibler (KL) Divergence profiles, information content (IC) scores, and nucleotide frequency matrices (motif blocks), yielding a 6W-dimensional vector (W = window width). All profile blocks were independently standardized before concatenation. t-SNE embedding was computed via consensus across 60 independent runs (perplexity = 15) to ensure robust genus-level clustering patterns. Genomes were classified by genus using hierarchical annotation rules (Table S1) applied to NCBI species nomenclature: exact species matches took priority, followed by epithet matching, then genus-level assignment. Ellipses in the visualization represent 2.5-sigma covariance contours for each genus/sub-genus cluster.

For all other analyses, which focused on *Leishmania* and *Trypanosoma*, genome assemblies and annotation files were sourced from TriTrypDB release 68 (May 7, 2024; https://tritrypdb.org/tritrypdb/app). For each species or strain, the upstream region flanking annotated start codons was analyzed, with the Kozak sequence defined as the 5 nucleotides immediately upstream of the start codon (positions -5 to -1). Only canonical ATG start codons were retained, and sequences were extracted from both strands. Statistical significance of nucleotide biases was assessed through chi-square analysis, comparing observed nucleotide frequencies to expected frequencies based on overall genomic GC content. The complete list of analyzed genomes includes: *L. panamensis* MHOM/COL/81/L13 (Lpan), *L. sp. Ghana* MHOM/GH/2012/GH5 (LspGhana), *L. sp. Namibia* MPRO/NA/1975/252LV425 (LspNamibia)*, L. tarentolae* Parrot-TarII-2019 (Ltar), *L. tropica* L590 (Ltrop), *L. turanica* LEM423 (Ltur), *L. aethiopica* L147 (Laeth), *L. amazonensis* PH8 (Lama), *L. arabica* LEM1108 (Larab), *L. braziliensis* MHOM/BR/75/M2903 (LbraM2903), *L. braziliensis* MHOM/BR/75/M2904 (LbraM2904), *L. donovani* BPK282A1 (LdBPK), *L. enriettii* MCAV/BR/2001/CUR178 (Lenri), *L. gerbilli* LEM452 (Lgerb), *L. infantum* JPCM5 (Linf), *L. major* Friedlin-2021 (Lmaj), *L. martiniquensis* MHOM/TH/2012/LSCM1 (Lmart), *L. mexicana* MHOM/GT/2001/U1103 (Lmex), *L. orientalis* MHOM/TH/2014/LSCM4 (Lori), *T. brucei* EATRO1125 (TbEATRO1125), *T. brucei* gambiense DAL972 (TbDAL972), *T. brucei* TREU927 (TbTREU927), *T. congolense* IL3000 (Tcongo), *T. cruzi* Berenice (TcBerenice), *T. cruzi* Brazil A4 (TcBrazilA4), *T. cruzi* Dm28c-2018 (TcDm28c), *T. cruzi* YC6 (TcYC6), *T. rangeli* SC58 (TrSC58). We also include the analysis of the genome of *L. donovani* Ld1S ^38^.

### Parasites

*Leishmania donovani* strain 1S (MHOM/SD/62/1S) was originally obtained from Henry Murray (Weill Cornell Medical College, New York, USA). All experiments were conducted *in vitro* using parasites with a low passage number (fewer than 10 passages from the primary hamster isolate stocks maintained in our laboratory ^37,38^). Promastigotes were axenically cultured in complete M199 medium (M199, 10% FBS, 20 mM HEPES; 100 μM adenine, 2 mM L-glutamine, 10 μg/ml folic acid, 13.7 μM hemin, 4.2 mM NaHCO_3_, 1xRPMI1640 vitamins, 8 μM 6-biopterin, 100 units penicillin and 100 μg/ml streptomycin, pH 7.4). Cultures were maintained at 26°C and routinely passed twice weekly by inoculating 1 × 10⁶ parasites into 5 mL of fresh medium. Growth kinetics were assessed by seeding an initial density of 1 × 10⁶ parasites/mL in 25 cm² cell culture flasks with complete medium, and total parasite numbers were quantified daily using a hemocytometer.

### Plasmid construction

The plasmid constructs were engineered to express an EGFP reporter gene under the control of distinct 5’ UTR sequences. Specifically, synthetic Kozak sequences (CCACC or CTTTA) were inserted immediately upstream of the EGFP start codon using different oligonucleotide primers (Supplementary Table S2). For high-throughput screening, a Kozak sequence library was generated using a degenerate primer (5’-NNNNN-ATG-EGFP-3’) to randomize the five nucleotides preceding the start codon. All EGFP expression sequences, each harboring a distinct Kozak sequence variant, were amplified using the Phusion™ High-Fidelity PCR Master Mix (New England Biolabs). The resulting PCR fragments and the linearized pLEXSY-neo vector, digested with *NotI* and *BamHI*, were assembled using Gibson Assembly Master Mix (New England Biolabs) in accordance with the manufacturer’s instructions. The assembled plasmids were introduced into 5-alpha Competent *E. coli* (New England Biolabs) by heat-shock transformation. Transformed cells were plated onto LB agar plates supplemented with 100 µg/mL ampicillin and incubated overnight at 37°C. Individual colonies were selected after 16 hours of growth for plasmid isolation. The integrity of all plasmid constructs and the fidelity of the inserted Kozak sequences were confirmed by Sanger sequencing (Eurofins Genomics). The plasmids were linearized prior to transfection using the SwaI restriction enzyme.

### Transfections and parasite selection

To assess the impact of Kozak sequence variants on EGFP reporter gene expression, *L. donovani* Ld1S promastigotes were transfected with linearized pLEXSY-neo2 (Jena Biosciences) constructs. 1 × 10⁷ promastigotes from mid-log culture phase were pelleted by centrifugation at 2,000 × g for 10 minutes at 4°C, resuspended in Tb-BSF buffer ^50^, and combined with 5 µg of each linearized, purified construct. Transfection was performed using an Amaxa Nucleofector™ 2b device (Lonza) with program X-001. Following the pulse, parasites were immediately transferred to a pre-warmed complete medium. Twenty-four hours post-transfection, stable transfectants were selected under 50 µg/mL Geneticin (G418; Sigma-Aldrich) pressure and subsequently cloned by limiting dilution in 96-well plates.

#### PCR

Genomic DNA was isolated from *L. donovani* promastigote harvested at mid-log culture phase. Briefly, a 1 mL aliquot of culture was centrifuged, and the resulting cell pellet was resuspended in 500 µL of DNAzol® reagent (Molecular Research Center, Inc.). Subsequent extraction steps, including the ethanol addition for precipitation, were carried out in accordance with the manufacturer’s standard protocol. The purified DNA pellet was washed, dried, and finally resuspended in nuclease-free water for downstream applications. Correct genomic integration of the pLEXSY construct at the *SSU* locus was confirmed by integration-specific PCR using one primer internal to the *neo* resistance cassette and a second primer annealing to the adjacent unmodified genomic region, as per the manufacturer’s (Jena Biosciences) recommendations (Supplementary Table S2).

### Western blot

1×10⁷ log-phase parasites were pelleted, washed with PBS, and lysed by boiling in 1× Laemmli buffer for 5 minutes. Proteins were separated by SDS-PAGE and transferred to a PVDF membrane. The membrane was probed with anti-GFP primary antibody (Santa Cruz, 1:2000) for 2 h, followed by HRP-conjugated goat anti-mouse secondary antibody (Santa Cruz, 1:4000) for 1 h. Blots were developed with Pierce™ ECL Western Blotting Substrate (Thermo Scientific™) and imaged on an Amersham™ ImageQuant™ 800.

### Flow cytometry

For EGFP fluorescence quantification, parasites were analyzed in a 96-well plate format using a CytoFLEX flow cytometer (Beckman Coulter). A minimum of 30,000 singlet events per sample were acquired, with live parasites gated based on forward-scatter (FSC) and side-scatter (SSC) profiles. For viability assessment, samples were stained with 2 µg/mL propidium iodide (PI, Sigma-Aldrich) for 15 minutes, and data were acquired using identical instrument settings. Data were analyzed and visualized using CytExpert software (v2.4.0.28). GFP fluorescence was detected in the GFP-A channel (exλ = 488 nm; emλ = 525/40 nm) and propidium iodide fluorescence was detected in the ECD-A channel (exλ = 488 nm; emλ = 610/20 nm) using the CytoFLEX flow cytometer (Beckman Coulter).

### Quantitative Reverse Transcription PCR (RT-qPCR)

1 × 10⁸ promastigotes from mid-log culture phase were pelleted by centrifugation at 2,000 × g for 10 minutes at 4°C. Total RNA was isolated using the NucleoSpin® RNA Plus kit (Macherey-Nagel) according to the manufacturer’s protocol, including an on-column DNase I digestion step to remove genomic DNA contamination. RNA concentration and purity were assessed spectrophotometrically. First-strand cDNA was synthesized from 1 µg of total RNA using M-MLV Reverse Transcriptase (Invitrogen) and oligo(dT) primers in a 20 µL reaction. Quantitative PCR (qPCR) was performed in 384-well plates using iTaq™ Universal SYBR® Green Supermix (Bio-Rad) on a LightCycler® 480 System II (Roche). Each 10 µL reaction contained 1 µL of diluted cDNA (equivalent to 100 ng of input RNA) and 100 nM of each gene-specific primer (sequences listed in Table S2). The thermocycling protocol consisted of an initial denaturation at 95°C for 1 min, followed by 40 cycles of 95°C for 10 s and 60°C for 30 s, with fluorescence data acquisition at the end of each 60°C step. A melt curve analysis was performed post-amplification to confirm primer specificity. Gene expression was normalized to two constitutive reference genes: *GAPDH* (Ld1S_300487200, LdBPK_303000.1, glyceraldehyde 3-phosphate dehydrogenase) and *DNA polymerase* (Ld1S_160127300, LdBPK_161640.1, DNA polymerase I). All primers used are shown in Supplementary Table S2. While for most analyses the primers were designed to recognize sequences in the coding region, for the pre-mRNA analysis the foward primer was designed to recognize a sequence in the 5’ UTR sequence present in the pre-mRNA but not in the mature mRNA, allowing us to differentiate between the two. Primer amplification efficiencies for each target were calculated from raw fluorescence data using the LinRegPCR version 2021.2 ^51^. Relative gene expression (fold change) was subsequently determined using the Relative Expression Software Tool (REST-MCS, version 2) ^52^. The software packages LinRegPCR and REST 2009 are publicly available at https://medischebiologie.nl/files/ and https://www.gene-quantification.de/download.html, respectively.

### Polysome profiling

Sucrose gradient ultracentrifugation and polysome fractionation were performed according to established methods ^53^. Briefly, promastigotes from mid-log culture phase were treated with 100 µg/mL cycloheximide (Sigma) for 10 min at 26°C. Cells were then harvested by centrifugation at 1,800 × g for 8 min at 4°C and washed once with ice-cold DPBS containing 100 µg/mL cycloheximide. After counting using a hemocytometer, aliquots of 5 × 10⁸ cells were pelleted, flash-frozen in liquid nitrogen, and stored at −80°C until lysis and sucrose gradient ultracentrifugation. Linear 10–50% sucrose gradients were prepared in SW 41 Ti rotor tubes using a gradient master 108 (BioComp). Lysates containing 15–20 A₂₆₀ units of polysomes were layered onto each gradient and centrifuged at 260,000 × g for 2 h at 4°C. Following centrifugation, gradients were fractionated using a piston gradient fractionator (BioComp) equipped with a UV detector, collecting 24 fractions of 500 µL each. Polysome profiles were monitored in real-time via UV absorbance from 254 to 260 nm. Collected fractions were processed immediately, and RNA was purified using TRIzol™ LS Reagent (Invitrogen) according to the manufacturer’s protocol. The only modification was the addition of glycogen (RNA grade) to improve pellet visualization and recovery. The samples were then analysed by RT-qPCR.

### Ribolace

To capture mRNA actively engaged in translation, ribosome-bound transcripts were isolated using the RiboLace Pro Kit (Immagina BioTechnology) ^54^, which employs affinity purification and magnetic separation. Promastigotes from mid-log culture phase were pelleted by centrifugation at 2,000 × g for 10 minutes at 4°C. The manufacturer’s standard protocol was followed with one modification: the nuclease digestion step was omitted to preserve full-length mRNA rather than ribosome-protected footprints.

### Drug Treatments

To assess mRNA stability and translation dynamics, *Leishmania donovani* promastigotes in mid-log culture phase were treated with the transcriptional inhibitor actinomycin D (Sigma, final concentration 10 µg/mL), or the translational inhibitor cycloheximide (Sigma, final concentration 30 µg/mL). Immediately before adding the drug (time 0) and at 3, 6, and 9 hours post-treatment, aliquots of parasites were collected by centrifugation. Cell pellets were flash-frozen in liquid nitrogen and stored at −80°C for subsequent total RNA extraction and mRNA analysis via RT-qPCR and/or RNA-seq.

### RNA Sequencing

For transcriptome analysis, total RNA was extracted from promastigotes at mid-log culture phase. Briefly, cells were pelleted by centrifugation at 2,000 × g for 10 minutes at 4°C and lysed in the buffer provided with the NucleoSpin® RNA Plus kit (Macherey-Nagel). All subsequent extraction steps, including on-column DNase I treatment, were performed according to the manufacturer’s protocol. RNA integrity was confirmed using an Agilent Bioanalyzer. Total RNA from all samples was used to generate strand-specific sequencing libraries with the Illumina Stranded mRNA Prep, Ligation kit. Briefly, polyadenylated mRNA was selectively captured from total RNA using magnetic beads, followed by enzymatic fragmentation. The fragmented RNA was then converted into double-stranded cDNA. During a subsequent ligation step, IDT for Illumina RNA UD Indexes Set A, Ligation was employed to attach unique dual-index (UDI) adapters to each sample, enabling sample multiplexing. A post-ligation cleanup was performed using Agencourt AMPure XP magnetic beads to purify the final libraries. Library quality and quantity were rigorously assessed prior to pooling and sequencing. Concentrations were measured fluorometrically using the Invitrogen Quant-iT Qubit dsDNA HS Assay Kit with Qubit Assay Tubes. Fragment size distribution and library integrity were analyzed on a Fragment Analyzer system using the High Sensitivity NGS kit (1–6000 bp) to confirm the absence of adapter dimers and validate the expected insert size range. Libraries were normalized based on Qubit and Fragment Analyzer data and pooled in equimolar ratios. The final pooled library was sequenced at the Biomics core facility (Institut Pasteur, Paris, France) on an Illumina NextSeq 2000 platform. Sequencing was performed in paired-end mode to generate high-depth, strand-specific transcriptome data for downstream bioinformatic analysis. Raw sequencing reads were processed using the Sequana RNA-seq pipeline (v0.18) ^49^. Adapters and low-quality bases were trimmed with fastp (v0.23.2). Cleaned reads were aligned to the *Leishmania donovani* strain 1S (Ld1S) genome using Bowtie2 (v2.5.4) with default parameters. Gene-level read counts were generated with featureCounts (v2.0.1; parameters: -t gene -g gene_id -s 2). Differential gene expression analysis was conducted in R (v4.1.2) using the DESeq2 package (v1.34.0). Normalization and dispersion estimation were performed with default DESeq2 parameters. Genes with an adjusted p-value (Benjamini-Hochberg correction) below 0.05 were considered statistically significant.

### GO enrichment analysis

GO enrichment analysis was performed to identify overrepresented biological processes within each gene set, using *Leishmania donovani* strain 1S (Ld1S) as the reference organism. The complete set of annotated genes from the Ld1S GFF files ^38^ served as the background population. Statistical significance for term enrichment was determined using a hypergeometric test, with a primary p-value cutoff of 0.05; Fisher’s exact test was also performed for comparative validation. To correct multiple hypothesis testing, the Benjamini-Hochberg False Discovery Rate (FDR) correction method was applied, and enriched GO terms were defined as those present in a query gene subset at a frequency significantly higher (FDR < 0.05) than expected by chance relative to the genomic background.

## RESULTS

### *Leishmania* spp shows a Kozak Signature distinct from other trypanosomatids

Kozak sequence profiles of the Euglenozoa genomes were characterized using the open-source KozakExplorer pipeline ^48^, which computes three complementary metrics (KL divergence, information content, and nucleotide frequencies) across a −20 to +6 position window relative to the AUG start codon (excluding the AUG itself). These feature vectors were then projected into two dimensions via consensus t-SNE (aggregated over 60 runs) to generate the embedding shown in Figure 1A. The result is a high-dimensional "fingerprint" of the translation initiation context across the analyzed genomes, which reveals a striking, non-overlapping segregation of kinetoplastid genomes according to their taxonomic genus. Notably, the embedding resolves not only the major genera *Trypanosoma* and *Leishmania*, but also clearly delineates the three *Leishmania* subgenera (*Leishmania, Viannia*, and *Mundinia*). This result indicates that the nucleotide context surrounding the start codon has undergone lineage-specific evolution, resulting in distinct translational initiation signatures. The robustness of these clusters is underscored by the consensus approach across 60 independent t-SNE runs, which mitigates stochastic variation and ensures that the observed grouping reflects intrinsic properties of the Kozak sequences.

**Figure 1.**
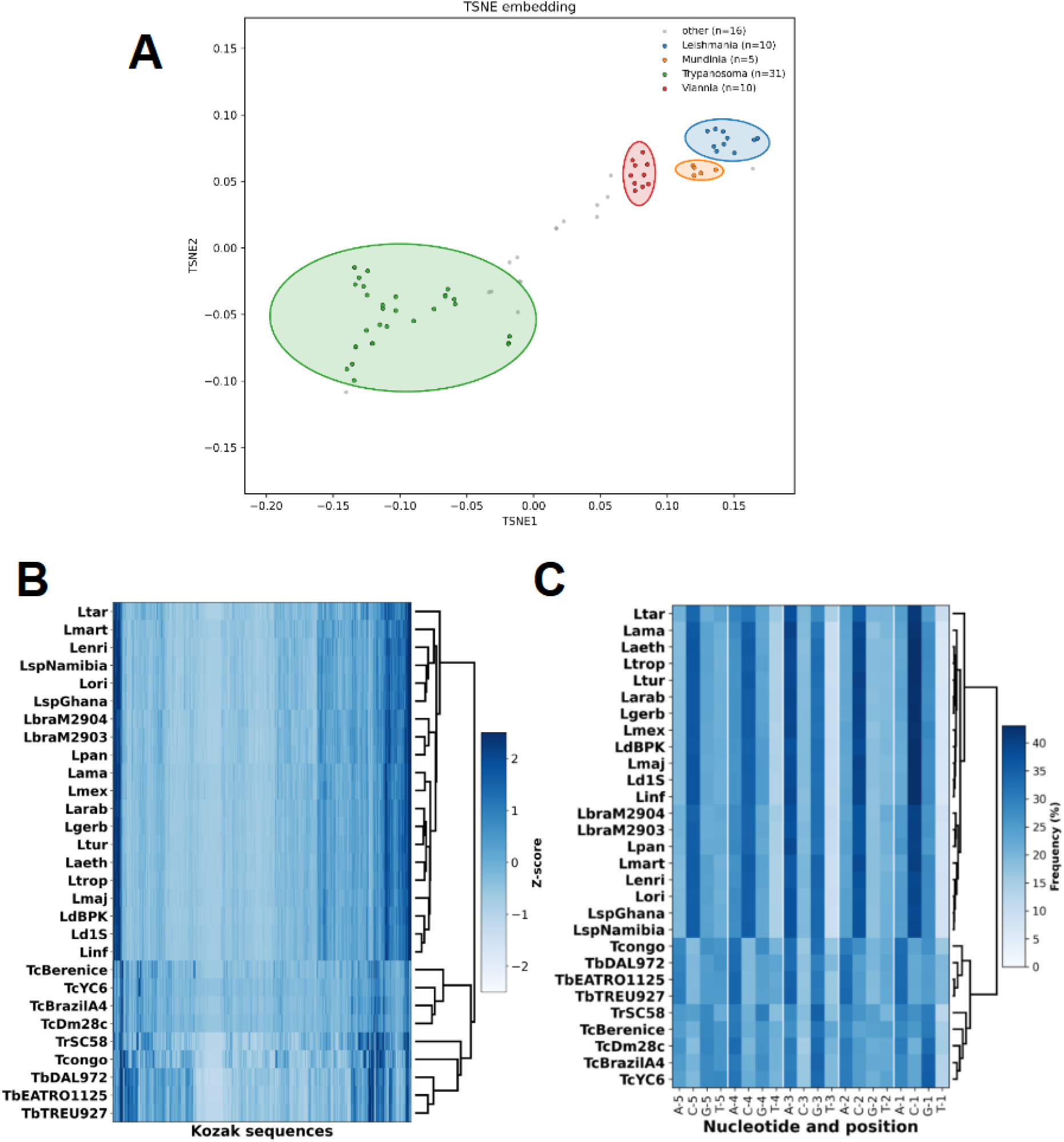
Landscape of Kozak sequence motifs in trypanosomatids. **(A)** t-SNE embedding of 72 Euglenozoa genomes. Genomes cluster into distinct groups corresponding to the major kinetoplastid genera or subgenera: *Trypanosoma* (n=31, green), *Leishmania (Leishmania)* (n=10, blue), *Leishmania (Viannia)* (n=10, red), *Leishmania (Mundinia)* (n=5, orange), and other euglenozoan lineages (n=16, gray). Ellipses highlight genus-level clusters. Embedding was computed from per-position Kozak profiles (KL divergence, information content, and nucleotide frequencies) via consensus t-SNE across 60 independent runs. **(B)** Z-score normalized heatmap showing the relative enrichment of Kozak sequences across Trypanosomatid species and strains. Dark blue indicates higher relative enrichment (sequences that have a higher percentage in that species than its own average across all species), while light blue/white indicates lower relative enrichment (sequences that have a lower percentage in that species than its own average across all species). Values with |Z| > 2 are considered statistically significant (p-value < 0.05). Kozak sequences are grouped based on similarity in their distribution patterns across species. Species are clustered based on similarity in their Kozak sequence preferences. **(C)** Nucleotide frequency analysis of Kozak sequences in trypanosomatids. The hierarchically clustered heatmap shows the frequency of each nucleotide at positions -5 to -1 upstream of the start codon across the indicated species and strains. The blue intensity represents nucleotide frequency, with darker blue indicating higher prevalence.

To define the Kozak sequence landscape in trypanosomatids, we also extracted the five-nucleotide region upstream of canonical ATG start codons (positions -5 to -1) from all protein-coding sequences (CDS) (Supplementary Table S3). The number of individual Kozak sequences detected per species ranged from 908 to 1,007 (Supplementary Figure S1A), indicating near-complete coverage of the theoretical repertoire (the theoretical space comprises 1,024 possible motifs considering 5 positions and 4 nucleotides).

The genome-wide Kozak sequence data were also visualized using two alternative heatmaps. Row-wise Z-score normalization (Figure 1B), which compares the relative abundance of each Kozak sequence across species, revealed striking differences between genera. *Leishmania* species displayed more homogeneous patterns, with Kozak sequences showing similar frequencies across all species. Significantly, clustering the heatmap according to frequency identifies distinct signatures at both the genus (*Trypanosoma* versus *Leishmania*) and subgenus levels *(Leishmania, Sauroleishmania, Mundinia,* and *Viannia*), similar to the consensus t-SNE analysis. This pattern suggests strong selection shaping the Kozak repertoire and supports an important role for this *cis*-regulatory element in the regulation of gene expression in trypanosomatids. Frequency analysis revealed a characteristic distribution: the majority of Kozak motifs are rare (i.e. low frequency, light blue), while a small subset of sequences is highly overrepresented across the genome (Supplementary Figure S1B). Maximum single-sequence frequencies varied substantially across species, from 0.61% to 3.98% (Supplementary Figure S1C). Remarkably, *T. cruzi* was the only species where any Kozak sequence exceeded 2% frequency, while *T. congo* showed the lowest maximum frequencies (0.61%). The dominant Kozak motifs in *T. cruzi* reflect the amplification of specific multi-gene families ^55,56^, likely due to duplication events that preserved not only the coding sequences but also their regulatory regions (Supplementary Table S4). This pattern is conserved across *T. cruzi* strains, as shown by analysis of the strains Dm28c, YC6, and BrazilA4. The GAGTG motif appears in 591, 567, and 442 CDSs, respectively, and is most frequently associated with Mucin-associated surface proteins (MASP) in all three strains. Similarly, the TCACG motif occurs 317, 466, and 277 times, respectively, and is predominantly linked to the mucin TcMUCII family. The TTTAT motif, showing 203, 288, and 268 occurrences, respectively, is primarily found in trans-sialidase genes (Supplementary Table S4).

Nucleotide frequency analysis at each position (Figure 1C) further highlighted these genus-specific signatures. *Leishmania* species presented a highly conserved CCRCC consensus motif and very low thymine frequency at positions -3 and -4. In contrast, *Trypanosoma* species showed a distinct pattern with elevated adenine frequency at positions -5, -4, -2, and -1, though this enrichment was less pronounced than the cytosine dominance observed in *Leishmania*, suggesting divergent translational initiation requirements between genera.

### Analysis of Initiation Codon Context in *L. donovani*

We next evaluated the Kozak sequence distribution in *L. donovani* Ld1S. From the 10,532 annotated sequences across the 36 contigs/chromosomes of the *L. donovani* LD1S genome, 8,491 protein-coding sequences (CDS) that annotate for a canonical ATG start codon were considered (Supplementary Table S3). A total of 930 distinct Kozak sequences were identified. The most common sequence, CAGCC, was observed 146 times, accounting for 1.72% of all occurrences in CDSs. In contrast, 124 Kozak sequences (or 13.3%) were singletons, each appearing only once in the dataset (Supplementary Figure S1).

In the 1980s, Marilyn Kozak suggested GCCGCCRCCAUGG as the optimal consensus sequence for the most efficient translation initiation. This sequence became known as the Kozak consensus, with specific positions being particularly crucial, especially a purine at the -3 position relative to the AUG ^57–61^. It is accepted that this consensus sequence has a similar effect in all eukaryotes, favouring the ribosome conformational change that terminates scanning of the mRNA molecule and initiates translation ^43,44^. Indeed, the *L. donovani* heatmap in Figure 2A shows an enrichment of the canonical CCACC motif at positions -5 to -1 relative to the start codon. However, it illustrates the per-nucleotide enrichment across the Kozak motif rather than reporting the frequency of the exact 5-nucleotide string CCACC. The CCRCC Kozak consensus motif, representing the two sequences CCACC or CCGCC, was identified in 155 CDS of the *L. donovani* Ld1S genome (1.8%). Despite the low frequency of the complete CCRCC motif, the conserved structure suggests selective pressure for efficient translation initiation in *L. donovani*. Statistical analysis revealed a significant bias (*p*<0.05) at positions -3, -2, and -1, with a notable enrichment for the ‘ACC’ motif preceding the ATG. Guanine (G) was moderately enriched at -3. Thymine (T) was consistently underrepresented across all positions, particularly at -1.

**Figure 2.**
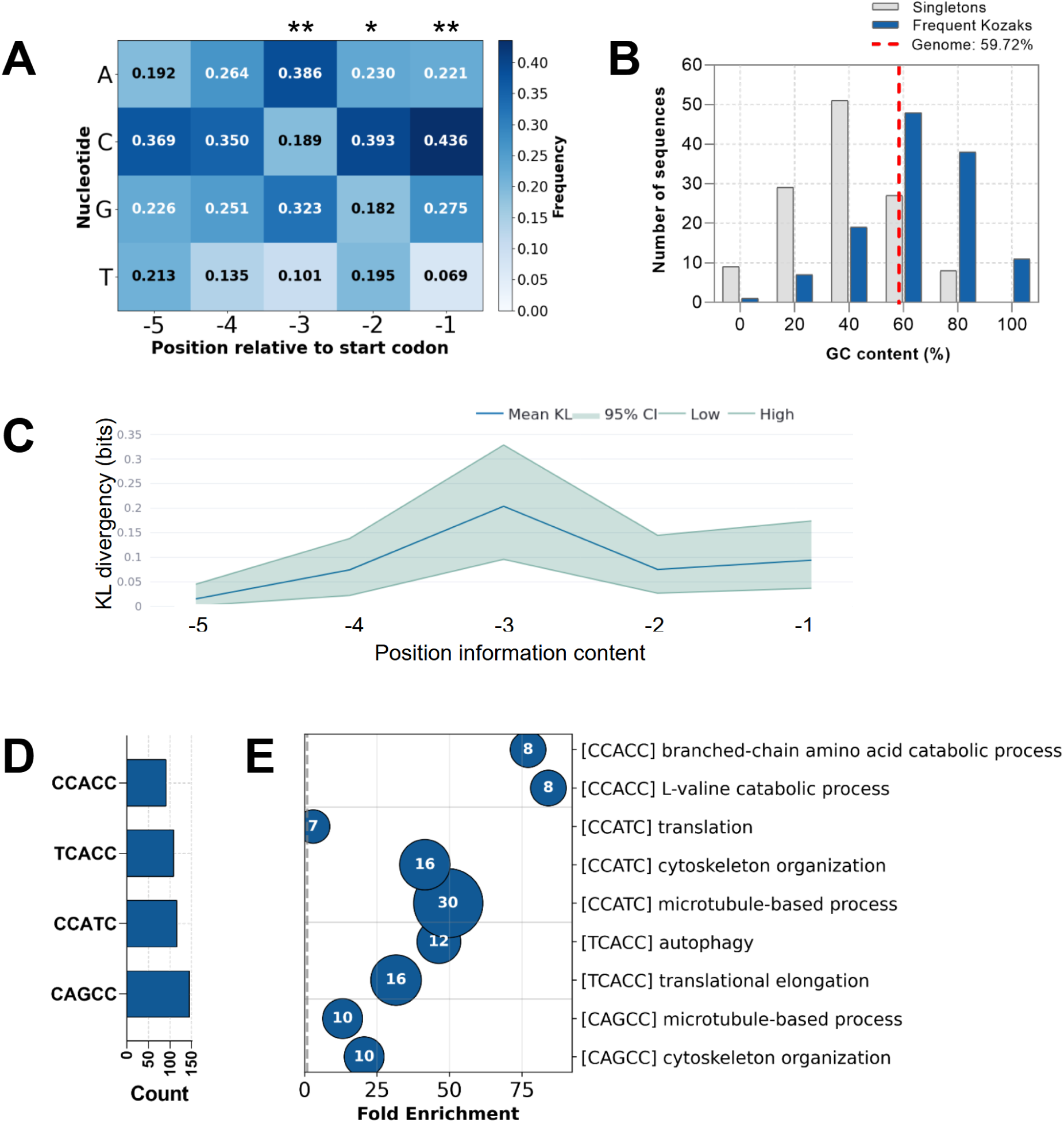
Genome-wide distribution of Kozak sequence motifs in *L. donovani*. **(A)** Heatmap showing nucleotide frequencies at positions -5 to -1 upstream of the start codon (AUG) for all coding sequences. Higher color intensity corresponds to higher frequency. Significance levels are indicated as follows: *, P ≤ 0.05; **, P ≤ 0.01 based on a Chi-square test. **(B)** GC content distribution of all 5-nt Kozak sequences (5-mers) in the 124 most frequent Kozak sequences and the "singleton" sequences (sequences occurring only once in *L. donovani* genome; n = 124). The vertical red line marks the overall genomic GC content of *L. donovani* (∼60%). **(C)** Per-site Kullback-Leibler (KL) divergence (bits) from the background nucleotide distribution, plotted against nucleotide position. **(D)** Frequency of the 4 most common 5-nt Kozak motifs. **(E)** Bubble plot displaying enriched Gene Ontology Biological Process terms for each of the 4 most common Kozak sequences. Terms shown are statistically significant with FDR-adjusted hypergeometric p-value < 0.05. The X-axis values show the fold enrichment value (observed/expected ratio), with a vertical dashed line at fold enrichment = 1 indicating no enrichment. Gene count is proportional to bubble size and displayed inside the bubbles.

To further investigate potential dependencies between adjacent nucleotides in the Kozak region, we performed a genome-wide analysis of dinucleotide frequencies in the 5 positions immediately upstream of start codons in *L. donovani*. Our analysis revealed significant non-random associations between neighboring bases (Supplementary Figure S2A), indicating that nucleotide preferences extend beyond single positions. CC at (-2, -1), CA at (-4, -3), and GC at (-3, -2) were the most enriched dinucleotides, and T-containing pairs were consistently under-represented across multiple positions. Similarly, analysis of trinucleotide frequencies across consecutive positions revealed distinct and position-dependent sequence preferences (Supplementary Figure S2B). At the upstream (-5, -4, -3) position, the highest frequencies were observed for CAG, TCA, CCA, GCA, CAA, and CGA. At the (-4, -3, -2) position, CAC emerged as the overwhelmingly dominant trinucleotide, followed by AGC and CGC. At the (-3, -2, -1) position immediately upstream of the reference site, GCC was the most frequent trinucleotide, accompanied by ACC and ATC. In contrast, stop codon-related triplets (TAA, TAG, TGA) and start codon triplet ATG were consistently depleted across all positions. These results demonstrate a clear, non-random gradient of di- or trinucleotide usage that is tightly coupled to positional context.

To investigate whether nucleotide composition influences Kozak sequence selection, we analyzed GC content distributions across two distinct categories of motifs: (1) all 124 "singleton" sequences that occur only once in the genome and (2) the 124 most frequent sequences (Figure 2B). This second category was matched in size to the 124 singletons to allow direct comparison. Comparative analysis revealed that high-frequency motifs are enriched for higher GC content, while rare, singleton motifs tend to be more AT-rich. High-frequency Kozak sequences (top 124 by occurrence) displayed a strong bias toward intermediate-to-high GC content. For this group of sequences, the 60% GC bin contained the largest proportion of sequences (38.7%, n=48), while the 80% bin was also well-represented (30.6%, n=38). Singleton sequences (n=124), in contrast, showed a markedly different distribution centered on lower GC content, with the distribution peaking sharply at 41.1% GC (n=51). Strikingly, no singleton sequences had 100% GC content, and the 60% and 80% GC bins were substantially less frequent (21.8% and 6.5%, respectively) compared with high-frequency Kozak sequences.

To quantify the information content at each Kozak position, we calculated the per-site Kullback-Leibler (KL) divergence from the background nucleotide distribution across all analyzed sequences (Figure 2C). This approach measures the expected log-ratio of probabilities under one distribution versus another, measuring how much information is lost when using the second distribution to approximate the first. Position -3 exhibited the highest KL divergence (mean = 0.208 bits), substantially exceeding all other positions (-5: 0.015-; 4: 0.071; -2: 0.079; -1: 0.105), indicating that this position is the most conserved and informative within the Kozak motif.

Gene Ontology enrichment analysis of transcripts containing the top four most frequent sequences — CAGCC (n = 146), CCATC (n = 116), TCACC (n = 109), and CCACC (n = 91) — revealed distinct functional associations (Figures 2D and 2E). For instance, the transcripts associated with CAGCC and CCATC motifs were enriched in cytoskeleton organization and microtubule-based processes, transcripts with CCACC were enriched in branched-chain amino acid catabolism, and CCATC and TCACC motifs were associated with transcripts linked to translation (Supplementary Table S5). In contrast, no significant enrichment was detected for the singleton group (n = 124) at an FDR < 0.05 (Supplementary Table S5).

Given the functional impact of Kozak sequences on translation efficiency and the observed Kozak preferences in *L. donovani*, we proceeded to experimentally investigate the "Kozak sequence code" and its role in regulating gene expression.

### Kozak Sequence Variants Modulate Gene Expression in *Leishmania*

To systematically assess the impact of various Kozak motifs on heterologous protein expression in *L. donovani*, we generated a pLEXSY-EGFP Kozak library for stable integration into the SSU ribosomal locus. This was achieved by using primers containing an NNNNN degenerate sequence that was added immediately upstream of the initiation codon (ATG) of EGFP during construct assembly (Supplementary Figure S3A). By randomizing the five positions upstream of the ATG we could obtain 1,024 possible nucleotide combinations. The resulting plasmid library was transfected into *L. donovani* promastigotes. Following drug selection, transfected parasites were cloned by limiting dilution into 96-well plates. EGFP fluorescence was quantified via plate reader for all wells during early log-phase growth (∼1 × 10⁶ parasites/mL). For the clones exhibiting fluorescence, genomic DNA was extracted, and the region encompassing the EGFP coding sequence and its upstream Kozak motif was amplified and analyzed by Sanger sequencing. Only wells containing a single, clearly defined Kozak sequence and clones without mutations in the EGFP coding region were included in the subsequent analysis. Data generated from clones with the same Kozak sequence were pooled to calculate standard deviations. Only Kozak sequences that are found more than once among the clones were analyzed. In total, 24 specific Kozak sequences represented by multiple clones were analyzed, showing various levels of EGFP fluorescence (Figure 3A), thus further confirming the *cis*-regulatory role of the Kozak sequence in *Leishmania*.

**Figure 3.**
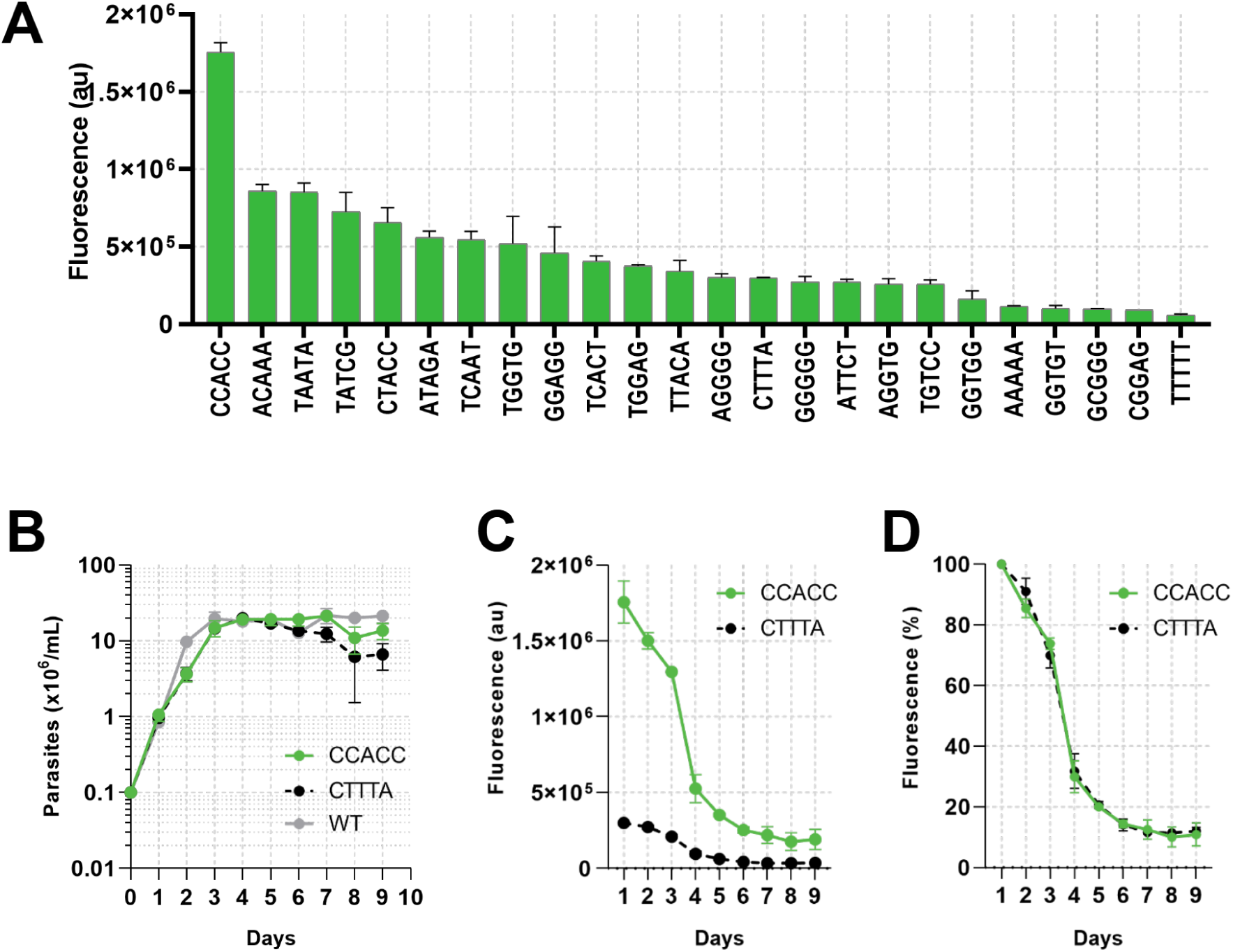
Modulating translation initiation efficiency in *Leishmania donovani* using a Kozak sequence library. **(A)** FACs analysis. A degenerate NNNNN region (-5 to -1) was cloned upstream of *egfp*, creating a library of variants. EGFP fluorescence was measured for individual clones during early log phase. Bars represent mean ± SD of all clones sharing the same Kozak sequence (data from multiple independent clones sharing the same motif were pooled for analysis). **(B)** Growth kinetics of WT and transgenic (CCACC-egfp, CTTTA-egfp) promastigotes *in vitro*. Cell density was determined by daily hemocytometer counts. Data represent mean ± SD (n=3 biological replicates). **(C)** Absolute fluorescence during growth was acquired by FACs (n=3 biological replicates). **(D)** Relative fluorescence during growth, with data normalized to day 1 levels of each strain (set to 100%), (n=3 biological replicates).

We next generated two pLEXSY constructs harboring EGFP under the control of either a canonical Kozak sequence (CCACC) or a rare, non-canonical variant (CTTTA), by using specific primers instead of degenerated ones. The CCACC motif occurs 91 times in the *L. donovani* genome and is associated with high translation initiation efficiency, whereas the CTTTA sequence appears only once and correlates with low mRNA abundance (Table S3). Following transfection and drug selection, integration of each construct into the small subunit (SSU) ribosomal RNA locus was validated by PCR (Supplementary Figure S3A and S3B). Amplification with primers annealing outside the integration site and within the EGFP coding sequence yielded the expected 1,983 bp product in transgenic clones, absent in wild-type or mock controls. We next evaluated the final protein output by Western blot analysis with an anti-EGFP antibody, which revealed that EGFP protein in the CTTTA strain was reduced to 17% of the level observed in the CCACC strain (Supplementary Figure S3C and S3D). This finding further validates the existence of a Kozak code in *Leishmania* and establishes a powerful experimental system for further mechanistic analysis.

No significant growth defect was observed in either transgenic line compared to the other (Figure 3B). We next monitored GFP levels of the different strains throughout parasite growth. Fluorescence analysis revealed that EGFP expression levels were strongly influenced by the Kozak sequence. Parasites carrying the CCACC Kozak sequence exhibited approximately six-fold higher GFP expression in early log-phase than those with the CTTTA variant during early log phase (Figure 3C). In both strains, GFP intensity decreased as cultures progressed through the growth curve. Strikingly, normalizing the GFP intensity to the initial value revealed identical fluorescence decay rates between the high- and low-expressing lines, despite their marked difference in absolute EGFP levels (Figure 3D). This observation indicates that the reduced expression in stationary parasites is not governed by translation initiation control, suggesting a possible reduction in the rate of transcription in stationary parasites. Similar patterns are observed at the transcript level for many genes in other trypanosomatids ^62^.

### Kozak-Dependent Translation Efficiency Influences mRNA Stability in *Leishmania*

To better understand how Kozak sequence variants influence gene expression, we compared EGFP mRNA abundance at different steps of the expression process in our transgenic *L. donovani* derivatives (CCACC vs. CTTTA).

First, we quantified EGFP pre-mRNA and mature mRNA levels using RT-qPCR on total RNA from early log-phase parasites. For the pre-mRNA analysis, we designed the foward primer to bind to a sequence of the 5’ UTR which is present in pre-mRNA but absent from mature mRNA, thereby allowing discrimination between the two. As anticipated, pre-mRNA levels were equivalent between the two strains, confirming that the Kozak sequence does not affect transcription initiation (Figure 4A). In contrast, mature EGFP mRNA levels in the CTTTA strain were only 34% of those in the CCACC strain (Figure 4B), suggesting a potential impact on mRNA stability or processing. To directly measure actively translated mRNA, we performed RT-qPCR on mRNA isolated via the RiboLace Pro Kit, which captures transcripts bound to actively translating ribosomes. The ribosome-associated EGFP mRNA in the CTTTA strain was also 34% of the CCACC level (Figure 4C).

**Figure 4.**
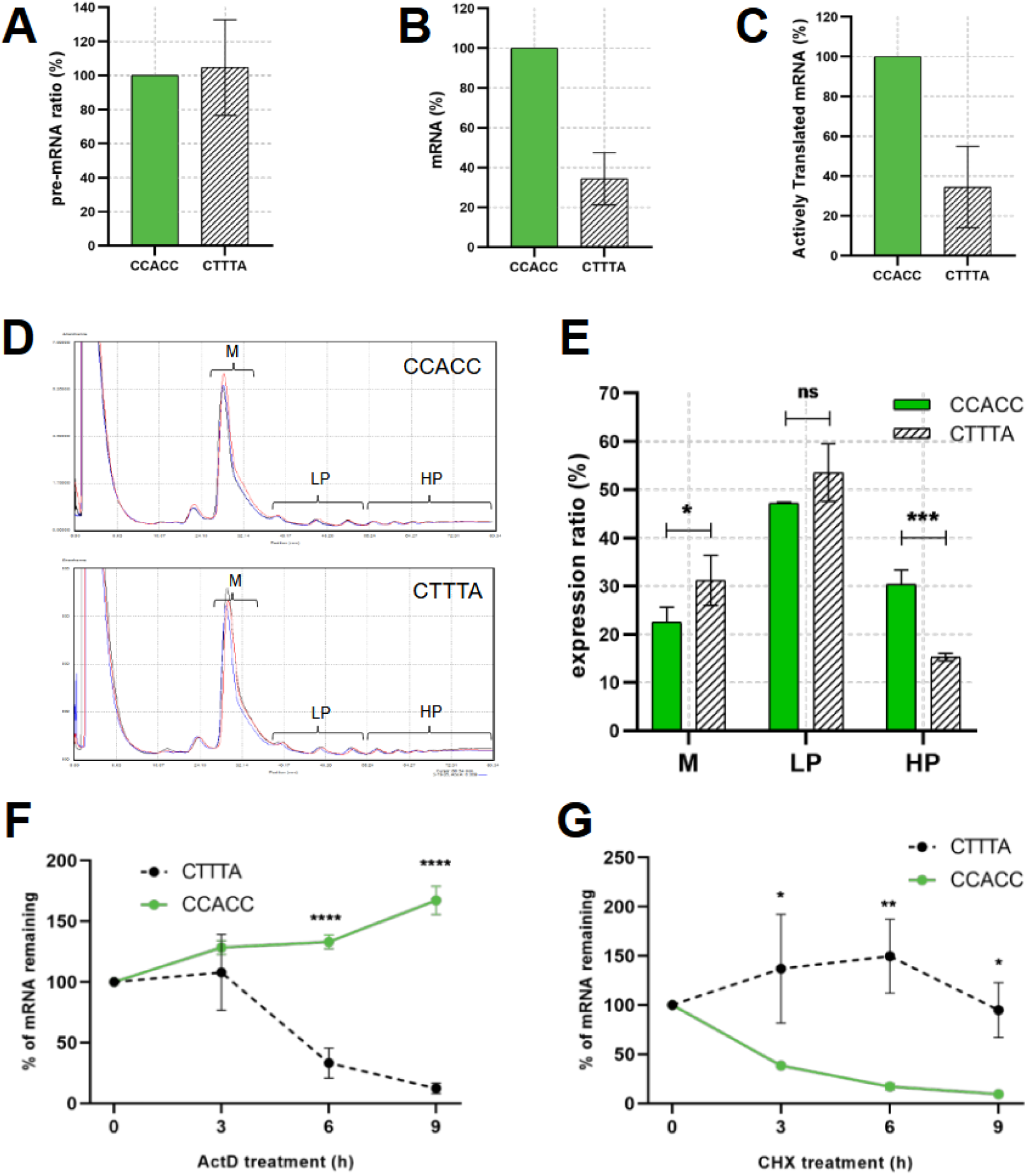
Mechanistic basis of Kozak sequence-mediated translation regulation in *L. donovani*. **(A)** Primary (pre-mRNA) transcript levels. **(B)** Steady-state mature mRNA levels. **(C)** Actively translating egfp mRNA associated with the ribosome fraction (recovered using RiboLace kit). For (A-C), data are normalized to *CCACC-egfp* levels (set to 100%). **(D)** Sucrose gradient polysome profiles for CTTTA and CCACC Kozak strains, indicating monosome (M), light polysome (LP), and heavy polysome (HP) fractions. **(E)** Relative abundance of EGFP mRNA in M, LP, and HP fractions, displayed as a percentage of total polysomal mRNA for each mutant. **(F-G)** egfp mRNA decay kinetics following inhibitor treatment: **(F)** Actinomycin D (10 µg/mL), and **(G)** Cycloheximide (30 µg/mL). All RT-qPCR experiment data are normalized to internal controls (*gapdh* and *dna polymerase*). Error bars indicate ± SD (n=3 biological replicates).

These results present a classic "chicken-versus-egg" scenario: either the inefficient translation initiation conferred by the weak Kozak sequence leads to mRNA destabilization, or the variant Kozak sequence compromises mRNA stability, which in turn limits the substrate for translation. To determine whether translation efficiency directly influences mRNA half-life in *Leishmania*, we next measured ribosome occupancy via polysome profiling, which yielded nearly identical profiles across both strains, confirming high experimental reproducibility and showing that the presence of eGFP with different Kozak sequences didn’t affect translation globally (Figure 4D). To quantify the distribution of EGFP mRNA across the polysome gradient by RT-qPCR, total EGFP mRNA in the unfractionated input was defined as 100% for each strain (Figure 4E). The proportion of EGFP mRNA in the monossomes (M), light polysome (LP), and heavy polysome (HP) fractions was expressed as a relative percentage of the total. LP samples contained 2-4 ribosomes on mRNAs. In the CTTTA-Kozak strain, a significantly greater proportion of EGFP mRNA was detected in the PP fraction, indicating lower ribosome occupancy. Conversely, in the CCACC-Kozak strain, EGFP transcripts were significantly enriched in the HP fraction, consistent with active translation by multiple ribosomes. These results confirm that the CCACC Kozak sequence supports a higher translation rate than the CTTTA variant.

To directly test whether the Kozak sequence variants CCACC and CTTTA influence mRNA stability, parasites expressing these sequences were treated with transcriptional and translational inhibitors using respectively actinomycin D (ActD) and cycloheximide (CHX). These treatments did not affect parasite viability during the 9-hour incubation period as shown by propidium iodide (PI) staining and flow cytometry (Supplementary Figure S4).

mRNA stability was assessed in ActD-treated parasites, which halts transcription and allows to monitor EGFP mRNA decay over time. This experiment revealed distinct degradation kinetics between the strains: EGFP mRNA in the CTTTA-Kozak strain decayed rapidly, whereas mRNA from the CCACC-Kozak strain was significantly more stable (Figure 4F). This inverse correlation between translation efficiency and mRNA turnover suggests that the efficiently translated CCACC-Kozak mRNA may be protected from degradation by its high ribosome occupancy, thereby enhancing its stability. On the other hand, translation-mediated mRNA decay is likely the main cause for reduction of CTTTA-Kozak mRNA abundance in untreated parasites. To test these possibilities, we next assessed the effect of the translation elongation inhibitor CHX on mRNA turnover. CHX-mediated translational inhibition had opposing effects on mRNA stability when compared to ACT treatment (Figure 4G): it destabilized CCACC-containing mRNA but stabilized CTTTA-containing mRNA, revealing translation-mediated mRNA decay as the prime cause for reduction of CTTTA-Kozak mRNA abundance in untreated parasites. This paradox pattern is reminiscent to published reports showing that the effect of CHX on both stabilizing or destabilizing specific transcripts depends on how these mRNAs are normally turned over ^61^.

### mRNA Stability Dynamics Following Inhibition of Transcription and Translation

To assess the role of translation-mediated decay on mRNA turnover at the genome-wide level, we treated wild-type parasites with ACT (10 µg/mL) or CHX (30 µg/mL) for 9 hours and performed RNA-seq analysis. Although this extended treatment likely induces substantial cellular stress (without affecting parasite viability, see Supplementary Figure S4), it provides the optimal condition to distinguish mRNA stabilizing and destabilizing effects, as validated by our EGFP reporter experiments (Figure 4).

Principal Component Analysis (PCA) was performed on variance-stabilized transformed (VST) normalized read counts from 12 samples (4 ACT-treated, 4 CHX-treated, and 4 non-treated (NT) controls). The tight clustering of biological replicates within each condition demonstrates good experimental reproducibility (Figure 5A). The first principal component (PC1), which explains 64.13% of the total variance, clearly separates the two treatment conditions, confirming that the treatment itself is the dominant source of variation. PC2 accounted for an additional 30.31% of the variance, resulting in a cumulative 94.44% of the variance explained by the first two components.

**Figure 5.**
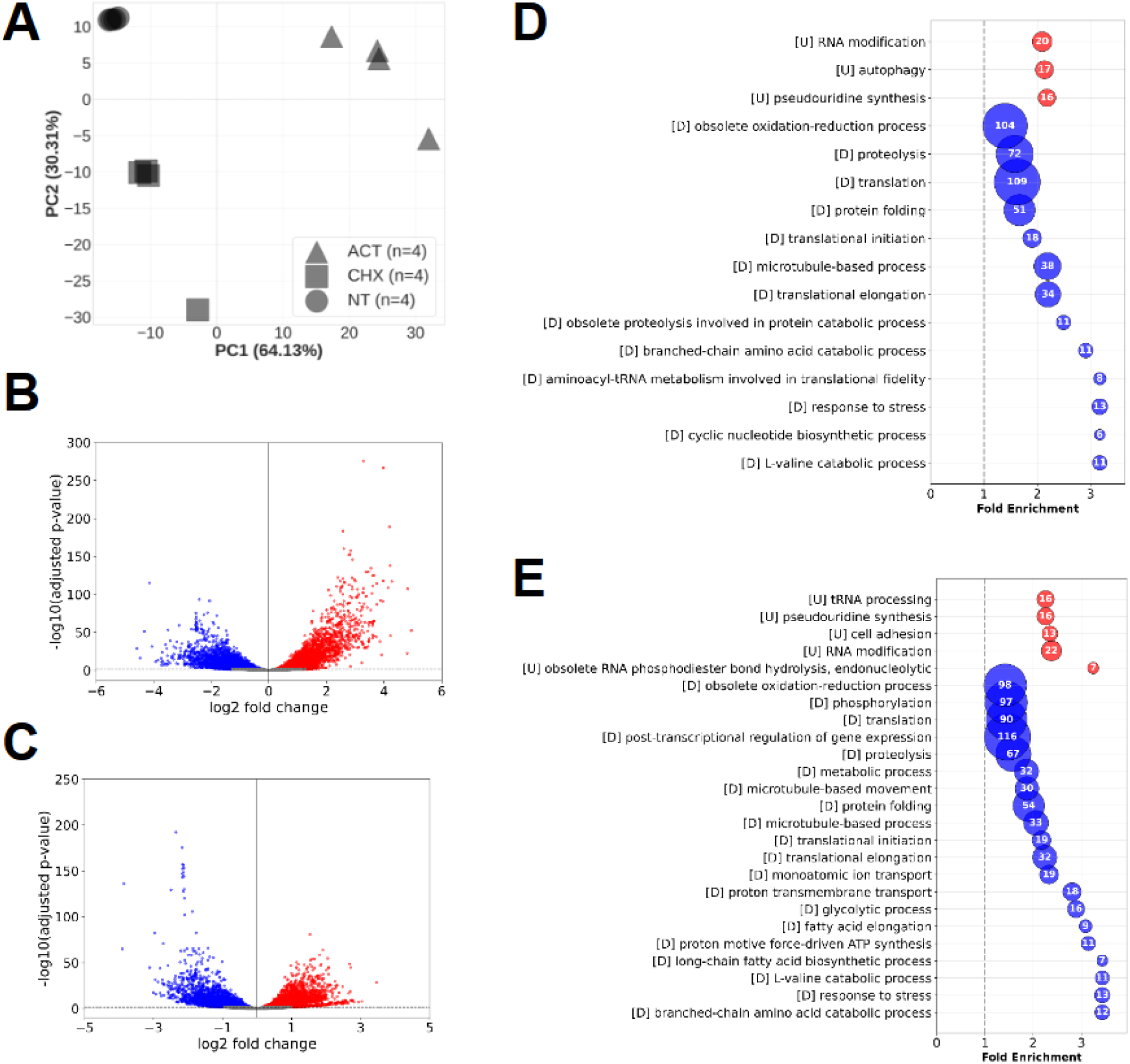
Global transcriptome changes in *L. donovani* following perturbation of transcription (ACT) and translation (CHX). Parasites were treated with Actinomycin D (10 µg/mL) or Cycloheximide (30 µg/mL) for 9 h. Non-treated parasites (NT) were used as control. The total RNA was extracted and sequenced. **(A)** Principal Component Analysis (PCA) of RNA-seq samples: Actinomycin D-treated (ACT), Cycloheximide-treated, and non-treated (NT). Points represent individual biological replicates (n=3 per condition), labelled by replicate ID (e.g., 1, 2, 3). Percentages indicate the variance explained by each principal component. **(B, C)** Volcano plots displaying differential gene expression following perturbation of transcription (ACT vs NT) **(B)** and translation (CHX vs NT) **(C)**. Each point represents an individual transcript. mRNAs with significantly increased (log₂FC > 0, padj < 0.05) and decreased abundance (log₂FC < 0, padj < 0.05) are highlighted in red and blue, respectively. **(D, E)** Bubble plot displaying enriched Gene Ontology terms for the category Biological Process for transcripts with significantly increased (red) and decreased abundance (blue) in ACT-treated **(D)** or CHX-treated **(E)** parasites versus NT control. Terms shown are statistically significant with FDR-adjusted hypergeometric p-value < 0.05. The x-axis shows the fold enrichment value (observed/expected ratio), with a vertical dashed line at Fold enrichment = 1 indicating no enrichment. Gene count is displayed inside the bubbles, and the bubble size is proportional to gene count. Bubble colour indicates regulation direction (red, enriched among the upregulated genes; blue, enriched among downregulated genes).

Analysis of differential gene expression between ACT-treated and control (NT) conditions revealed that, from the 10,532 detected transcripts, 6,773 showed significant differential expression (padj < 0.05). Among these, 3,400 transcripts showed increased abundance, while it was decreased for 3,373 transcripts (Figure 5B and Supplementary Table S6). CHX treatment produced an even more pronounced effect, with 7,520 transcripts exhibiting significant differential expression, including 3,740 transcripts with increased, and 3,780 transcripts with decreased abundance (Figure 5C and Supplementary Table S6). Assessing Gene Ontology (GO) terms associated with the differential expression changes in response to the different inhibitors used in this study identified various core biological processes affected by both treatments, including RNA metabolism (processing, modification, snoRNA transcription), post-transcriptional regulation, phosphorylation, proteolysis, and translation (Figure 5D, 5E, and Supplementary Table S7).

### Kozak Sequence Preferences Define Co-Regulated and Counter-Regulated Transcripts with Distinct Functional Implications

We next exploited our RNA-seq data to assess translation-mediated mRNA decay at a whole transcriptome level. A double-ratio plot comparing expression changes under ACT and CHX treatments, both relative to non-treated controls (Figure 6A), places differentially expressed transcripts into four quadrants based on their response patterns (Supplementary Table S6). Overall, three global expression groups can be distinguished in this double ratio plot. The first group of 6,007 transcripts did not show statistically significant differences in abundance in one of the two condition and thus were excluded from further analysis. A second group includes transcripts that show the same expression changes in response to both treatments, including 1,417 transcripts with increased (Quadrant (Q) 1) and 1,574 transcripts with decreased abundance (Q3), reflecting common responses likely induced by the stress imposed by both ‘expression’ inhibitors. The third and most interesting group includes transcripts that are showing opposite expression changes in response to ACT and CHX, just as observed for in our EGFP reporter strains (see Figure 4), i.e., increased abundance in response to ACT but decreased abundance in response to CHX (Q2, 805 transcripts) and *vice versa* decreased abundance in response to ACT but increased abundance in response to CHX (Q4, 729 transcripts). Gene Ontology enrichment analysis (Supplementary Table S7) associated Q2 with autophagy, protein heterodimerization and nucleosome localization (Figure 6B), whereas Q4 showed most significant enrichment in cell adhesion and DNA-binding transcription factor activity (Figure 6C).

**Figure 6.**
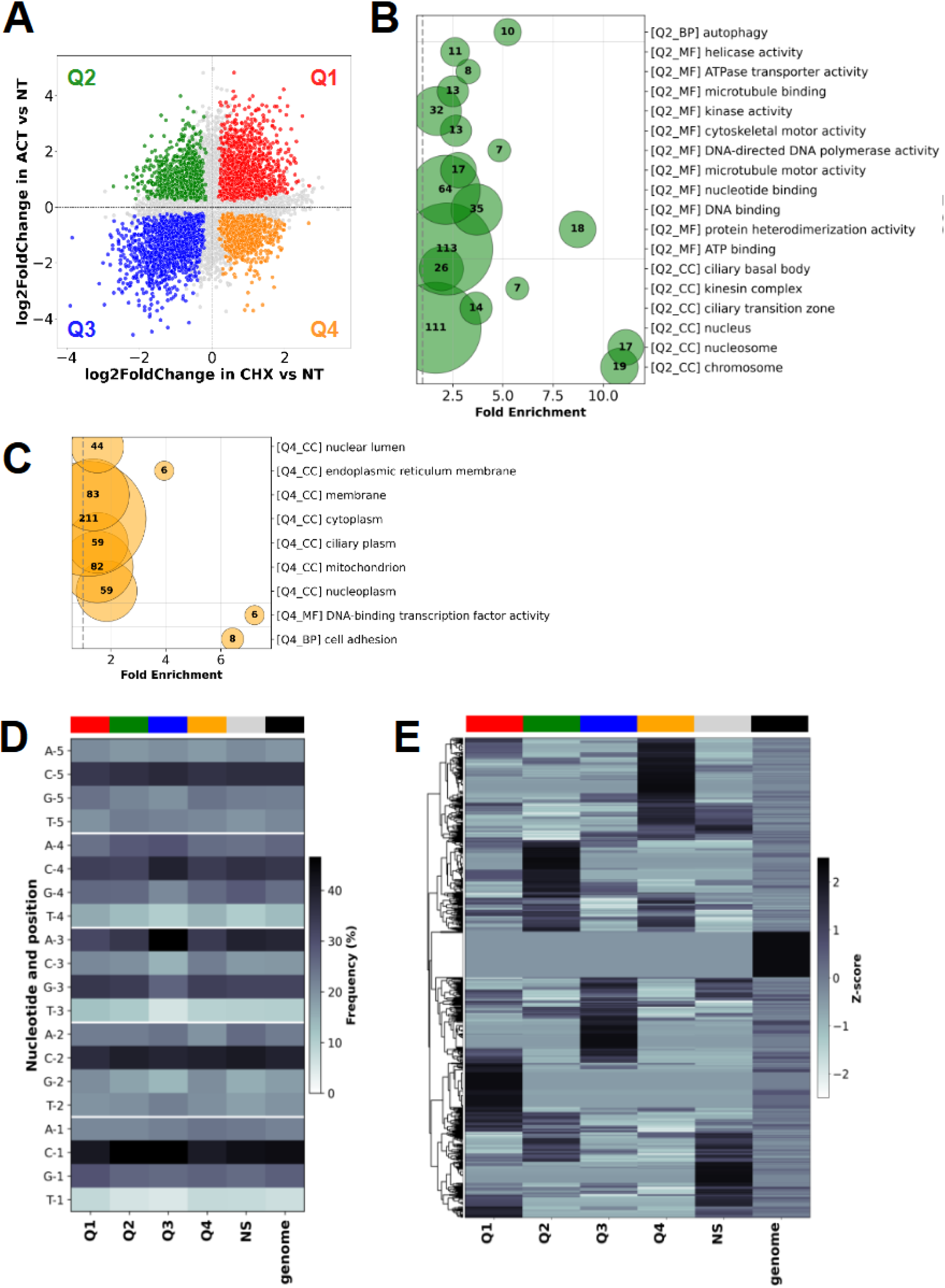
**(A)** Scatter plot showing log₂ fold change values for ACT versus NT on the Y-axis and CHX versus NT on the X-axis. Colours denote transcripts with significant differential expression (adjusted p-value < 0.05): red for upregulated in both treatments, blue for downregulated in both treatments, green for upregulated in ACT and downregulated in CHX, and orange for downregulated in ACT and upregulated in CHX. Gray points represent non-significant transcripts. **(B-C)** Bubble plot displaying enriched Gene Ontology terms for transcripts falling into quadrants Q2 **(B)** and Q4 **(C)** of the double-ratio plot. GO terms are categorized into three ontologies: Molecular Function (MF), Cellular Component (CC), and Biological Process (BP). Terms shown are statistically significant with FDR-adjusted hypergeometric p-value < 0.05. The X-axis values show the fold enrichment value (observed/expected ratio), with a vertical dashed line at Fold enrichment = 1 indicating no enrichment. The bubble size is proportional to gene count (number of genes from the quadrant associated with that GO term), with the numerical value indicated inside the bubble. The bubble plots are coloured according to their corresponding quadrant of the double ratio plot shown in panel (A). **(D)** Nucleotide frequency analysis of Kozak sequences in the four double-ratio plot quadrants, the non-significant (NS) group of transcripts, and the Ld1S genomic background. The heatmap shows the frequency of each nucleotide at positions -5 to -1 upstream of the start codon across all sets. The red-colored heat map represents nucleotide frequency, with darker red indicating higher prevalence. **(E)** Z-score-normalized heatmap of Kozak sequence enrichment across the four double-ratio plot quadrants, the Ld1S genomic background, and the non-significant (NS) group of transcripts. Dark red indicates higher relative enrichment (sequences that have a higher percentage in that set than its own average across all sets), while light red/white indicates lower relative enrichment (sequences that have a lower percentage in that set than its own average). Values with |Z| > 2 are considered statistically significant (p-value < 0.05). Kozak sequences (Y axis) are grouped based on similarity in their distribution patterns across sets. No clustering was done for the different sets (X axis).

We next assessed if the separation of the transcripts into four quadrants based on their response pattern to the two inhibitors correlated with distinct Kozak motifs, which would further corroborate their role in expression regulation (Figure 6D). Consistent with our previous analysis (Figure 2C), plotting the nucleotide frequency of Kozak sequences across the four quadrants revealed nucleotide position -3 as the most informative as it showed the highest variability thus differentiating between quadrants. Assessing the distribution of Kozak motifs across the four quadrants revealed Q3 as having both the highest number of distinct Kozak variants and the highest frequency of certain motifs (Supplementary Figures S5A–C). Notably, the canonical motif CCRCC was present in the top 5 five of the most common Kozak motifs in nearly all sets, with Q1 being the sole exception (Supplementary Figure S5D). A more contrasting signal emerged when assessing Kozak sequence distributions by plotting their Z-score-normalized frequencies across the four double-ratio plot quadrants (Figure 6E). The resulting heatmap identifies distinct Kozak Z-score profiles that are specific for each quadrant, suggesting that the Kozak motif is not merely a translational enhancer but also a *cis*-acting determinant of mRNA fate, likely through differential recruitment of trans-acting factors that couple translation elongation to mRNA decay ^64^. This observation reinforces the notion that Kozak-mediated patterns may constitute a regulatory layer, potentially defining Kozak-governed regulons in *Leishmania*.

## DISCUSSION

In this study, we combined bioinformatic, genetic, and pharmacological approaches to investigate the impact of Kozak sequences on *Leishmania* gene expression. We identified the presence of a conserved Kozak consensus in *Leishmania* characterized by selective sequence constraints that seem to have evolved to fine-tune translation initiation in a species-specific manner. Using EGFP reporter strains, we provide initial evidence for Kozak-mediated, translation-dependent regulation of mRNA turnover in *Leishmania*, thus revealing a previously unrecognized layer of regulation. Our results shift the prevailing paradigm for gene regulation in *Leishmania* occurring mainly at the 3’UTR and underscore the necessity of integrating studies of the 5’UTR into future models of parasite adaptive responses. Our findings open a series of important questions (i) on the evolution of trypanosomatid Kozak sequences and their role in species-specific translational control, (ii) on the *Leishmania* pathways that link translation initiation and ribosome stalling to mRNA decay, and (iii) on the existence of a putative Kozak code that allows for co-regulation of functionally linked genes and pathways.

Seminal work by Lukes et al. (2006) ^46^ and Stanton & Mensa-Wilmot (2006) ^47^ demonstrated that nucleotides at positions -1 to -3 upstream of the AUG initiation codon modulate protein synthesis in *Leishmania*, and notably, that a purine at position -3 is critical for efficient translation. Despite these foundational findings, in-depth mechanistic analyses on Kozak-mediated regulation in these parasites were still missing. Our results expand on these earlier findings. First, our observation that Kozak-like sequences have diverged in a lineage-specific manner between *Leishmania* and *Trypanosoma* supports their biological relevance as functional, cis-regulatory elements. Despite the close relationship of these parasites with respect to gene content and synteny ^65^, only *Leishmania* preserves a conserved canonical CCRCC Kozak motif, providing additional proof of its important function in these pathogens. In contrast, we did not identify such a conserved Kozak motif in *Trypanosoma*, confirming a previous study in *T. brucei*, which instead revealed a strong Kozak context among a subpopulation of constitutively expressed transcripts ^66^. Such inter-species divergence reveals different Kozak evolutionary trajectories, which may be driven by co-evolution of species-specific translation initiation machineries ^44,67,68^. Indeed, trypanosomatids are characterized by expansion of various protein families interacting with mRNA, including the translation initiation factors eIF4E and eIF4G or the RNA-binding protein (RBP) PABP ^69^. This expansion likely enables selective mRNA translation, as distinct combinations of these *trans*-acting factors may recognize specific *cis*-acting UTR motifs ^70–74^. Although *Trypanosoma* and *Leishmania* share expansion of these protein families, they differ in the actual numbers of paralogs, their regulation and possible activity ^69,74–77^. Conceivably, this difference may have affected the selective pressures acting on the Kozak motif, with RBPs and translation initiation factors in trypanosomes offering alternative initiation control mechanisms, enabling a more relaxed Kozak evolution compared to *Leishmania*.

A second key finding that endows the *Leishmania* Kozak sequence with important biological function is its direct link to mRNA stability. In model organisms, translation initiation is a major determinant of mRNA turnover: decay rates are inversely related to initiation efficiency, and perturbations that reduce initiation accelerate deadenylation and decapping ^78–80^. Likewise, several studies in trypanosomatids indicate that translation governs mRNA turnover through initiation efficiency, elongation dynamics, and codon-driven ribosome occupancy ^31,81,82^. Using EGFP reporter strains as a robust experimental framework for dissecting Kozak function, we obtained direct evidence for their implication in controlling translation initiation-dependent mRNA stability. Upon ACT treatment, strongly expressed CCACC-containing EGFP mRNAs showed slower decay rates than weakly expressed CTTTA-containing EGFP mRNAs. CHX treatment rescued the unstable transcripts from decay, directly linking translatability to transcript stability and demonstrating that Kozak-mediated differences in initiation efficiency drive differential mRNA turnover in *Leishmania*. Surprisingly, an opposing effect was observed for highly translated CCACC-containing EGFP mRNAs, suggesting a decay pathway that acts on transcripts with stalled ribosomes. These observations are reminiscent to previous studies in yeast, where transcriptional changes were observed in specific mRNA subsets following CHX treatment ^83^, or in Drosophila ^78^, where transcripts with weak Kozak sequences were largely insensitive to fluctuations in elongation rates but highly sensitive to changes in global initiation rates, while mRNAs with strong Kozak sequences show the opposite pattern. Just like proposed for Drosophila, the differential sensitivity to ACT and CHX observed for distinct mRNA pools in *Leishmania* may provide a mechanistic basis for how global variations in translation rates, whether acting on initiation or elongation, can selectively reshape the proteome by modulating transcript output. Rather than being detrimental, sub-optimal Kozak sequences likely provide an additional layer of post-transcriptional regulation, fine-tuning translational responses to cellular stress or environmental changes ^64,78^.

The opposing effect of the inhibitors on different mRNA pools informs on distinct mRNA decay mechanisms in *Leishmania* acting at different stages of the translation cycle. Low initiation rates and reduced ribosome occupancy favour recruitment of deadenylases, such as CCR4–NOT components conserved in trypanosomatids ^84^, which trigger decapping followed by 5′→3′ exonucleolytic degradation ^63,85^. It has further been proposed that the stability of any given transcript depends on a competition between the eIF4F initiation complex and the decapping complex for access to the 5’ methylguanosine cap ^80^. Unlike most eukaryotes, where decapping is mediated by the conserved Dcp1/Dcp2 complex ^86,87^, this process is catalyzed in trypanosomatids by the ApaH-like phosphatase ALPH1 ^88^. CHX, by arresting elongation and stalling ribosomes on the mRNA, impedes access of the decapping/5′→3′ decay machinery thereby stabilizing initiation-limited transcripts, as observed for the weak-Kozak EGFP reporter. In contrast, high initiation rates and increased ribosome occupancy efficiently protects mRNAs against decay. As observed in other eukaryotes, CHX-induced ribosome stalling in *Leishmania* on these strongly translated mRNAs may be sensed by the DEAD-box helicase Dhh1/DDX6, again conserved in trypanosomatids ^89^, promoting decay by interacting with the deadenylation complex CCR4–NOT as described above. Even though a non-canonical, nonsense-mediated mRNA decay (NMD) pathway has been identified in trypanosomatids ^90,91^, its implication in the observed inhibitor-mediated rescue or decay of Kozak variant EGFP transcripts seems unlikely, given that NMD is triggered by premature termination codons or abnormally long 3′ UTRs rather than distinct Kozak motifs ^92,93^. Likewise, alternative forms of mRNA degradation in *Leishmania* not initiated by deadenylation or decapping are depending on 3′ UTR elements ^94^ and thus can be ruled out in Kozak-mediated translational control.

Finally, our data suggest that Kozak sequences may participate in the coordinated regulation of distinct gene sets in *Leishmania*, as evidenced by the divergent Kozak profiles associated with expression changes in ACT- and CHX-treated parasites. Such a Kozak code has been linked to co-regulation of gene clusters in other organisms, including *T. brucei, Cryptococcus* spp., Drosophila and Zebrafish (*Danio rerio*) ^66,78,95–97^. Conceivable, this *cis*-acting regulatory code requires co-evolution of a matching interface at the level of *trans*-acting factors or ribosomal structures, both of which have been observed. For example, in *Saccharomyces cerevisiae,* specific Kozak sequences with adenosine at the −3 position facilitate translation via interaction with RPS26, while absence of this factor causes ribosomes to bind and translate a distinct set of poorly translated mRNAs, often linked to stress responses ^64^. Furthermore, structural differences between rabbit and yeast ribosomes and initiation factors have been linked to different preferences in start codon sequence contexts ^98^. The possibility of specialized ribosomes that may impact on mRNA translatability and subsequently mRNA stability in trypanosomatids is strongly supported by stage-specific and fitness-adapted differences in snoRNA expression, rRNA modification or the differential expression of ribosomal proteins ^35,99–101^. Furthermore, *Leishmania* stage differentiation has been recently linked to quantitative and qualitative changes in ribosomal components, implicating stage-specific protein kinases in the establishment of specialized ribosomes ^37^.

In conclusion, our results draw a complex picture of translational regulation in *Leishmania* that is governed by *cis*-acting regulatory motifs inside 5’ and 3’UTRs in interaction with co-evolved trans-acting factors implicated in RNA-binding or translation initiation. In the absence of canonical, promoter-driven transcriptional control ^7,27,102^, *Leishmania* may have accentuated such translational gene-expression control, exploiting Kozak-mediated differences in initiation efficiency and mRNA stability to fine-tune protein production across changing environmental conditions. Such a regulatory interface at the translational level would further provide a powerful biological filter to harness the intrinsic instability of the *Leishmania* genome for adaptation and fitness gain, by avoiding the phenotypic expression of toxic gene dosage effects, while promoting beneficial ones. Dissociating the molecular mechanisms underlying such a filter is a major challenge for future studies and will require (i) a comprehensive investigation of 5′ UTR-mediated regulation through long-read RNA sequencing, CLIP experiments (cross-linking and immunoprecipitation) or CRISPR-based perturbation assays that will respectively reveal UTR isoforms and upstream open reading frames (uORFs) ^96,103–108^, identify trans-acting factors recognizing specific Kozak motifs, and functionally validate regulatory interactions. Progress will also depend on improved genome annotation, which is critical for accurately defining translation initiation sites and, consequently, Kozak contexts and other regulatory motifs. Finally, the limited availability of Gene Ontology annotations for a substantial fraction of genes largely constrains the identification of functional regulons and broader regulatory networks, whose expression may be co-ordinated at translational levels ^109^.

## DATA AVAILABILITY

RNAseq raw data were submitted to the European Nucleotide Archive (ENA, https://www.ebi.ac.uk/ena/browser/home), ArrayExpress accession E-MTAB-16907. The data is not publicly released yet but can be requested during the review process.

## SUPPLEMENTARY DATA

### Tables

Table S1 – t-SNE Table S2 – Primers

Table S3 – Kozaks_trypanosomatids Table S4 – Frequent_Kozaks_Tcruzi

Table S5 – Kozaks_Ld1S_genome_GOterms Table S6 – RNAseq_Ld1S_ACTvsCHXvsNT

Table S7 – RNAseq_Ld1S_ACTvsCHXvsNT_GOterms

### Supplementary figure legends

**Supplementary Figure S1 – Expanded dataset of Kozak sequence motifs in trypanosomatids. (A)** Diversity of Kozak sequence repertoires across Trypanosomatid species. The plot displays the number of unique Kozak sequences detected in each species (from a maximum of 1024 possible motifs considering 5 positions and 4 nucleotides). **(B)** Distribution of Kozak sequence frequencies across Trypanosomatid species. Each data point represents a single Kozak sequence and indicates its percentage frequency within each genome, calculated relative to the total number of gene Ids of that genome. **(C)** Maximum Kozak sequence frequency by species. The bar plot displays the highest percentage frequency achieved by a given Kozak sequence within each Trypanosomatid species. Each bar is annotated with the specific Kozak sequence(s) achieving this maximum frequency.

**Supplementary Figure S2 – Expanded dataset of Kozak contexts in the *L. donovani* genome. (A)** Position-specific dinucleotide enrichment. Color intensity indicates the number of occurrences of each dinucleotide at adjacent positions, evaluated across four consecutive two-position windows: (−5, −4), (−4, −3), (−3, −2), and (−2, −1). Statistical significance was assessed using a chi-square test across these four windows. **(B)** Trinucleotide enrichment across position triplets. Color intensity shows the number of occurrences of each 3-mer at consecutive positions. Trinucleotides are arranged in four columns with each 16 rows for layout purposes only. Statistical significance was determined using a chi-square test applied to the full set of 64 trinucleotides across three consecutive three-position windows: (-5, -4, -3), (-4, -3, -2), and (-3, -2, -1). Significance levels are indicated as follows: ns, P > 0.05; *, P ≤ 0.05; **, P ≤ 0.01; ***, P ≤ 0.001; ****, P ≤ 0.0001.

**Supplementary Figure S3 – Cloning strategy of the Kozak sequences and effect on GFP expression. (A)** Schematic of the pLEXSY-EGFP integration strategy. Variable Kozak sequences (NNNNN) were inserted upstream of the *egfp* coding sequence. Green arrows indicate primers used for PCR validation. The dotted line marks the chromosomal integration site in the parasite genome. **(B)** PCR validation of transgenic parasite lines. A ∼2 kb product confirms site-specific integration for mutants with CCACC or CTTTA Kozak sequences, but not in wild-type (WT) parasites or the no-template negative control (NC). MW, molecular weight marker. **(C)** Western blot for phenotypic characterization of ccaccEGFP and ctttaEGFP strains. The left panel shows a Western blot using an anti-GFP antibody; the right panel shows the corresponding Coomassie-stained gel for normalization purposes. **(D)** Bar graph showing quantified protein levels based on densitometric analysis of Western blot (WB) signals, with the CCACC-driven EGFP signal set to 100%.

**Supplementary Figure S4 – Cell viability assessment**. Representative flow cytometry histograms of PI fluorescence, showing the PI positive control (heat-killed parasites, 5 min at 70°C, black line), and the parasites treated for 9 hours with ACT (red line) or CHX (orange line).

**Supplementary Figure S5 – Expanded dataset of Kozak contexts in the RNA-seq.** Kozak sequence analysis encompassed the four double-ratio quadrants, NS sequences, and the Ld1S genomic background. **(A)** The plot displays the number of total gene IDs in each set and the number of unique Kozak sequences detected in each set (Kozak sequences were counted only once per set, regardless of how many gene IDs they were associated with, so that repeated occurrences of the same sequence were not double-counted). **(B-C)** Distribution of the Kozak sequence in counts **(B)** and in frequencies relative to the number of gene IDs in each set **(C)**. Each individual data point represents a different Kozak sequence. **(D)** Frequency of the 5 most common 5-nt Kozak motifs in each set.

## ACKNOWLEDGEMENTS

We thank Peter Myler for the generous support and insightful guidance throughout this project. We thank Institut Pasteur Biomics Platform C2RT for technical assistance.

## AUTHOR CONTRIBUTIONS STATEMENT

AMMS: Conceptualization, Data curation, Formal analysis, Investigation, Methodology, Project administration, Validation, Visualization, Writing – original draft, Writing – review & editing

TC: Conceptualization, Data curation, Formal analysis, Investigation, Methodology, Software, Validation, Visualization, Writing – review & editing

JPL: Investigation, Methodology, Visualization, Writing – review & editing PJ: Investigation, Methodology, Writing – review & editing

ET: Investigation, Methodology, Visualization, Writing – review & editing CCRA: Investigation, Methodology, Visualization, Writing – review & editing

ALK: Investigation, Methodology, Resources, Supervision, Writing – review & editing ZNK: Investigation, Methodology, Resources, Supervision, Writing – review & editing

GFS: Conceptualization, Funding acquisition, Project administration, Resources, Supervision, Writing – original draft, Writing – review & editing

