## Supplementary figures and images for "Initiation codon context governs translation-coupled mRNA decay and coordinated expression in the human parasite *Leishmania*"

### Supplementary data

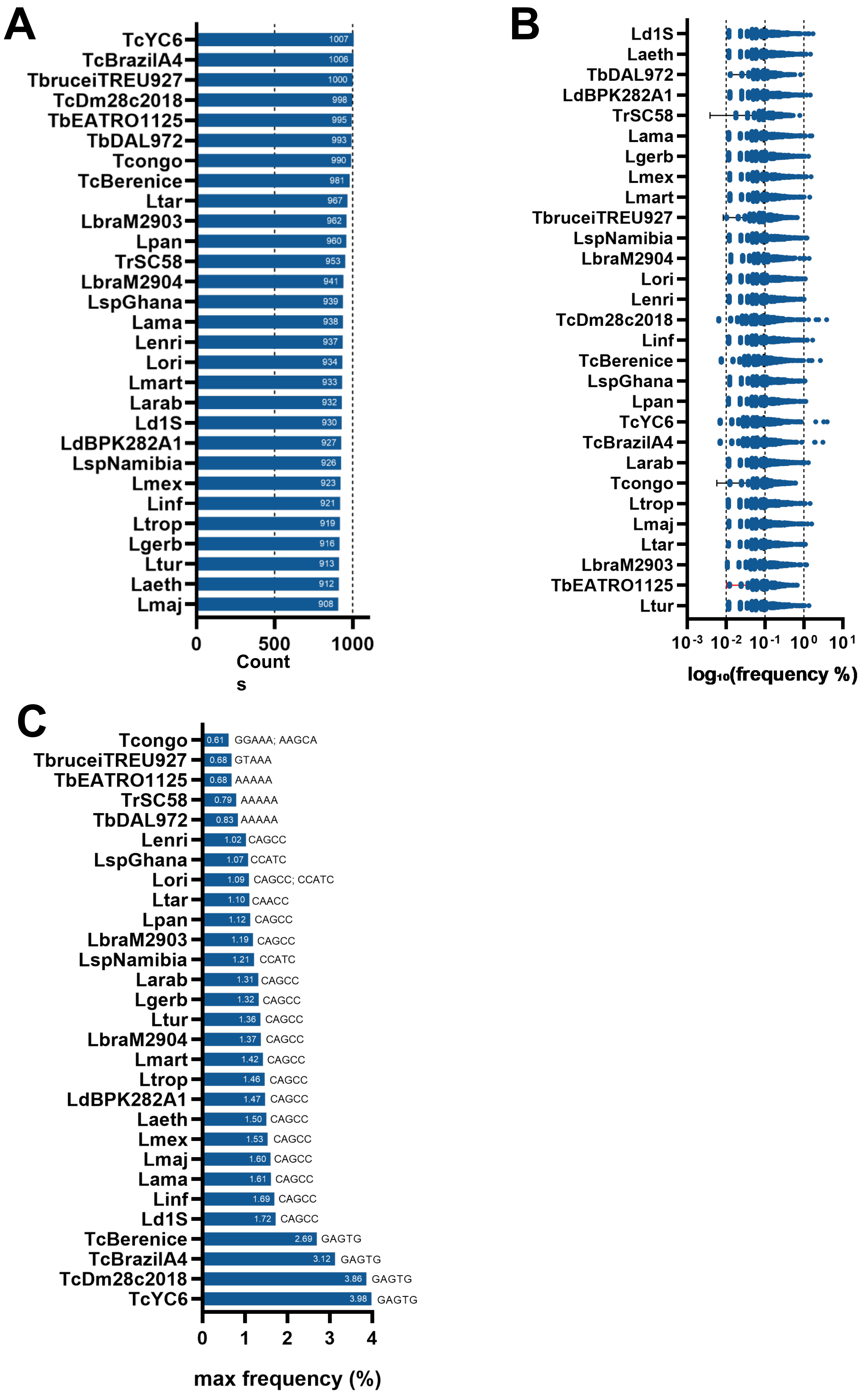

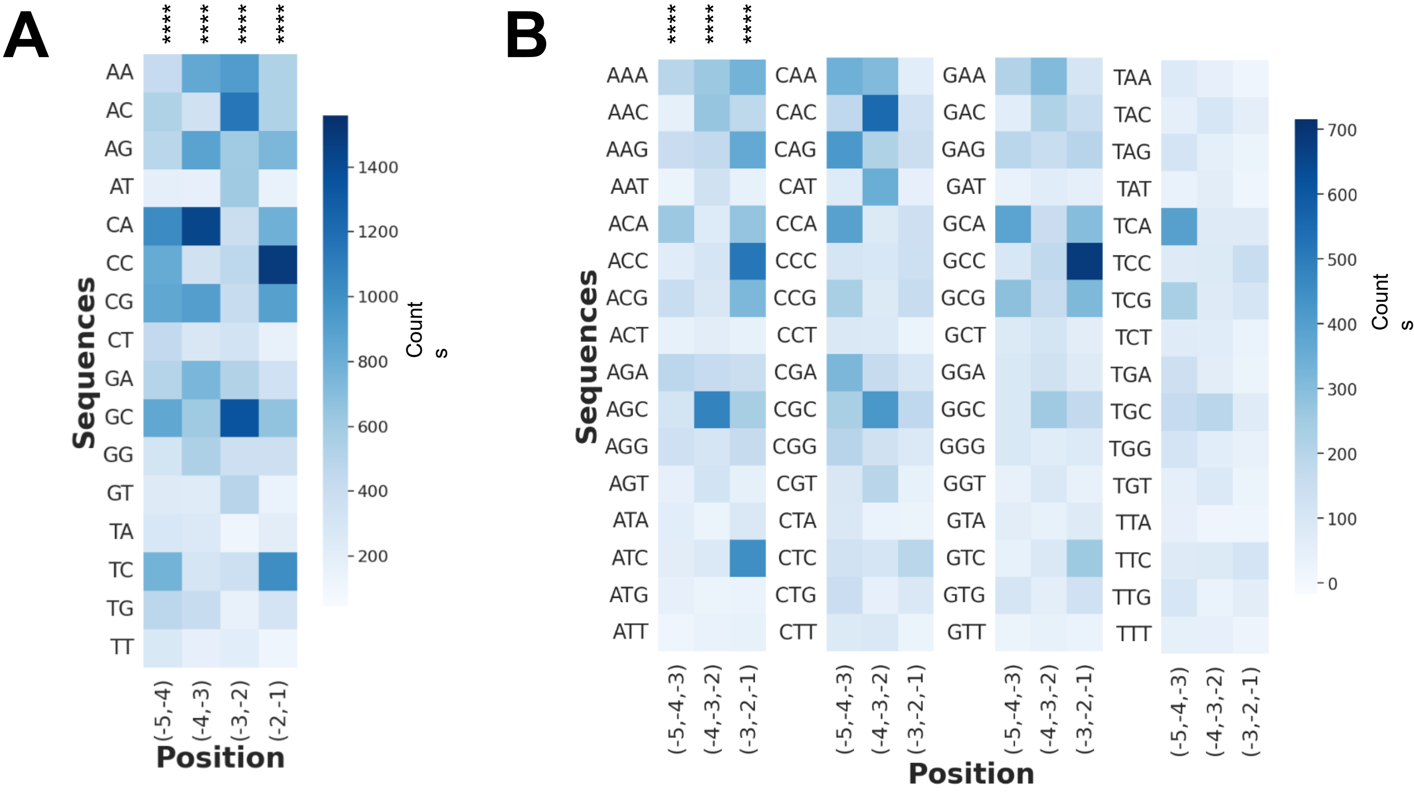


**Figure S2**

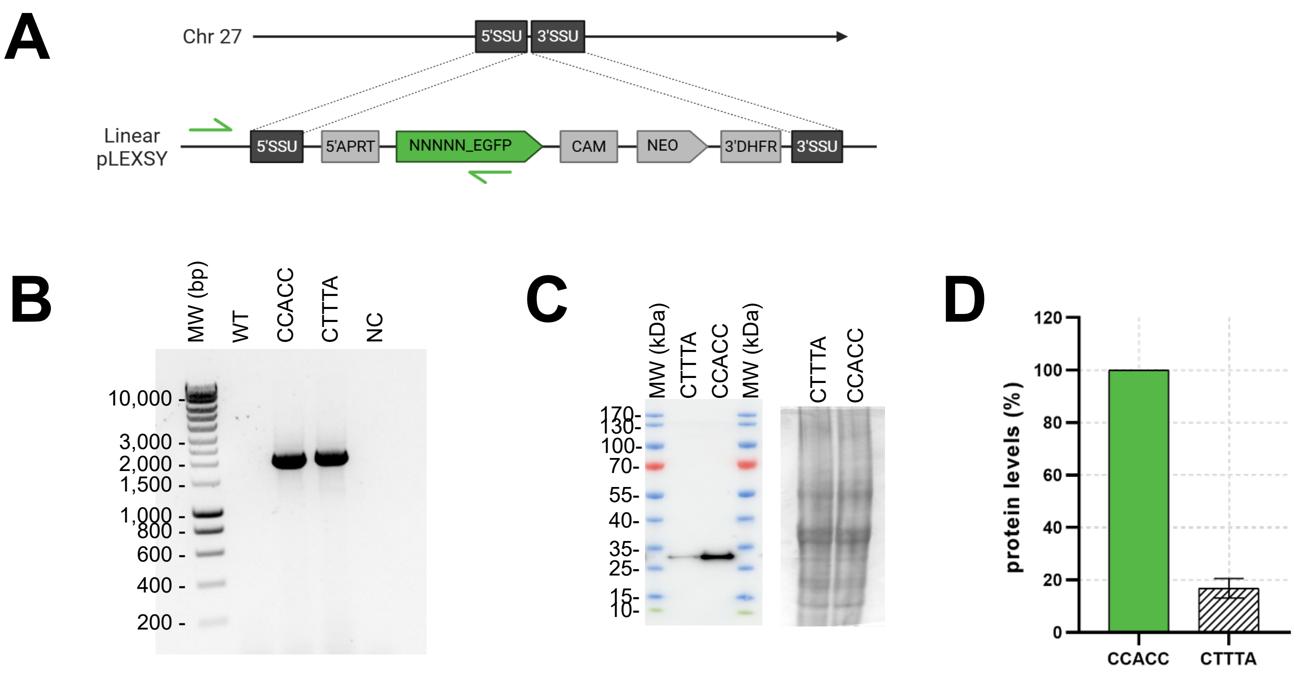

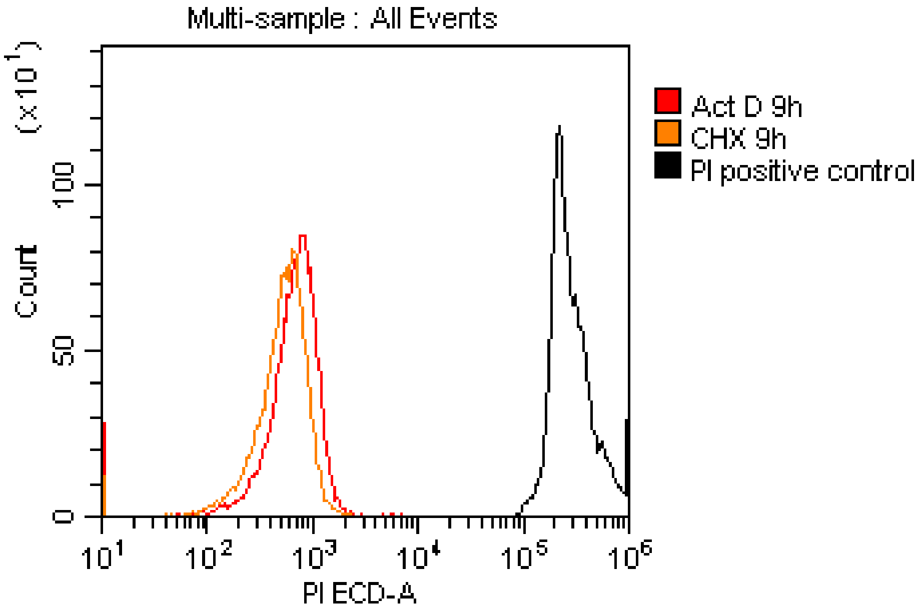

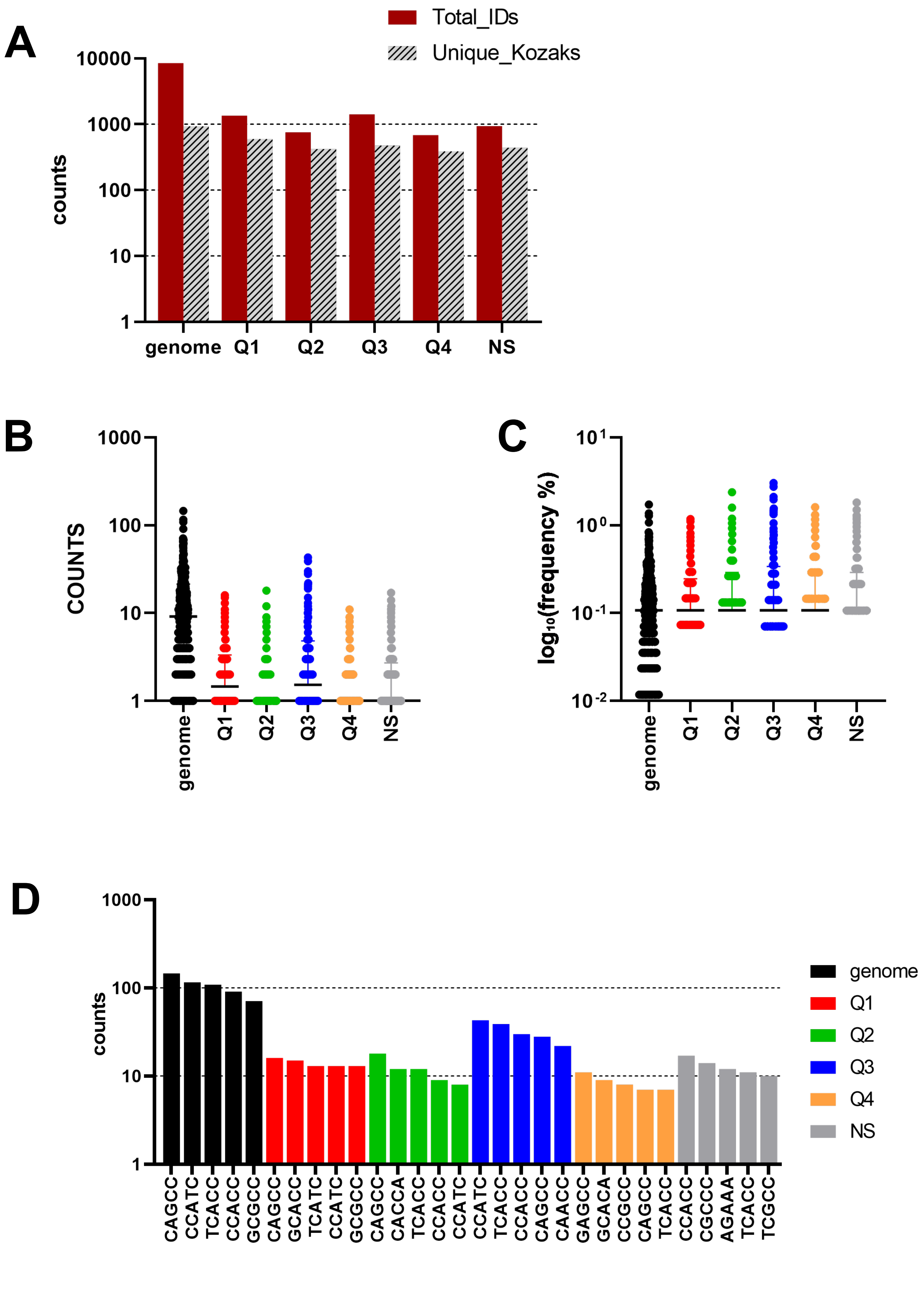
